# Epigenetic aging of human hematopoietic cells is not accelerated upon transplantation into mice

**DOI:** 10.1101/271817

**Authors:** Joana Frobel, Susann Rahmig, Julia Franzen, Claudia Waskow, Wolfgang Wagner

## Abstract

Transplantation of human hematopoietic stem cells into immunodeficient mice provides a powerful *in vivo* model system to gain functional insights into hematopoietic differentiation. So far, it remains unclear if epigenetic changes of normal human hematopoiesis are recapitulated upon engraftment into such “humanized mice”. Mice have a much shorter life expectancy than men, and therefore we hypothesized that the xenogeneic environment might greatly accelerate the epigenetic clock. We demonstrate that genome-wide DNA methylation patterns of normal human hematopoietic development are indeed recapitulated upon engraftment in mice – particularly those of normal early B cell progenitor cells. Furthermore, we tested three epigenetic aging signatures and none of them indicated that the murine environment accelerated age-associated DNA methylation changes. These results demonstrate that the murine transplantation model overall recapitulates epigenetic changes of human hematopoietic development, whereas epigenetic aging seems to occur cell intrinsically.

## Introduction

Humanized mice (HuMice) are used for a wide variety of applications in biomedical research, ranging from tumor biology, over studies of human hematopoiesis, to vaccine testing (Shultz et al., 2012; Willinger et al., 2011). Within the last decades various mouse models have been generated to improve hematopoietic reconstitution. For example, KIT-deficient NOD/SCID *Il2rg*^*−/−*^ *Kit*^*W41/W41*^ (NSGW41) mice support stable engraftment of lymphoid and myeloid cells without the need for irradiation conditioning prior to transplantation, allowing analysis of human hematopoietic cells in a steady-state condition (Mende et al., 2015; Rahmig et al., 2016). Phenotypically, humanized mice reflect multilineage differentiation that closely resembles human counterparts. However, it was yet unclear if transplanted human cells recapitulate epigenetic changes of normal hematopoietic development. Furthermore, mice have a significantly shorter life-span than men and this might result in faster epigenetic aging upon transplantation into the faster aging cellular environment (Wagner, 2017). In this study, we have therefore analyzed global DNA methylation (DNAm) profiles of human hematopoietic cells that stably engrafted in mice.

## Results and Discussion

Hematopoietic stem and progenitor cells (CD34^+^) were isolated from human umbilical cord blood (CB) and transplanted into five NSGW41 mice (Cosgun et al., 2014). Nineteen weeks after transplantation, the bone marrow (BM) was harvested and flow cytometric analysis revealed that 96.4% ± 1.9% of hematopoietic cells were of human origin. Immunophenotypic analysis of these human CD45^+^ (hCD45^+^) cells reflected differentiation toward lymphoid (B cells, T cells, and NK cells) and myeloid lineages (monocytes, granulocytes, and immature granulocytes; Fig. 1A). The majority of the engrafted human cells expressed CD19 and therefore seemed to be committed toward B cell development (71% ± 3%; Fig. 1B). We analyzed genome-wide DNAm patterns of sorted hCD45^+^ cells with Infinium HumanMethylation450 BeadChips. In comparison to DNAm profiles of various mature human hematopoietic subsets (GSE35069) (Reinius et al., 2012), unsupervised hierarchical clustering (Fig. 1C) and principle component analysis (PCA; Fig. 1D) demonstrated that epigenetic profiles of HuMice were overall still closely related to CD34^+^ CB cells (GSE40799) (Weidner et al., 2013). This was somewhat unexpected, because the engrafted cells clearly reflect immunophenotypic changes of hematopoietic differentiation.

**Fig. 1.**
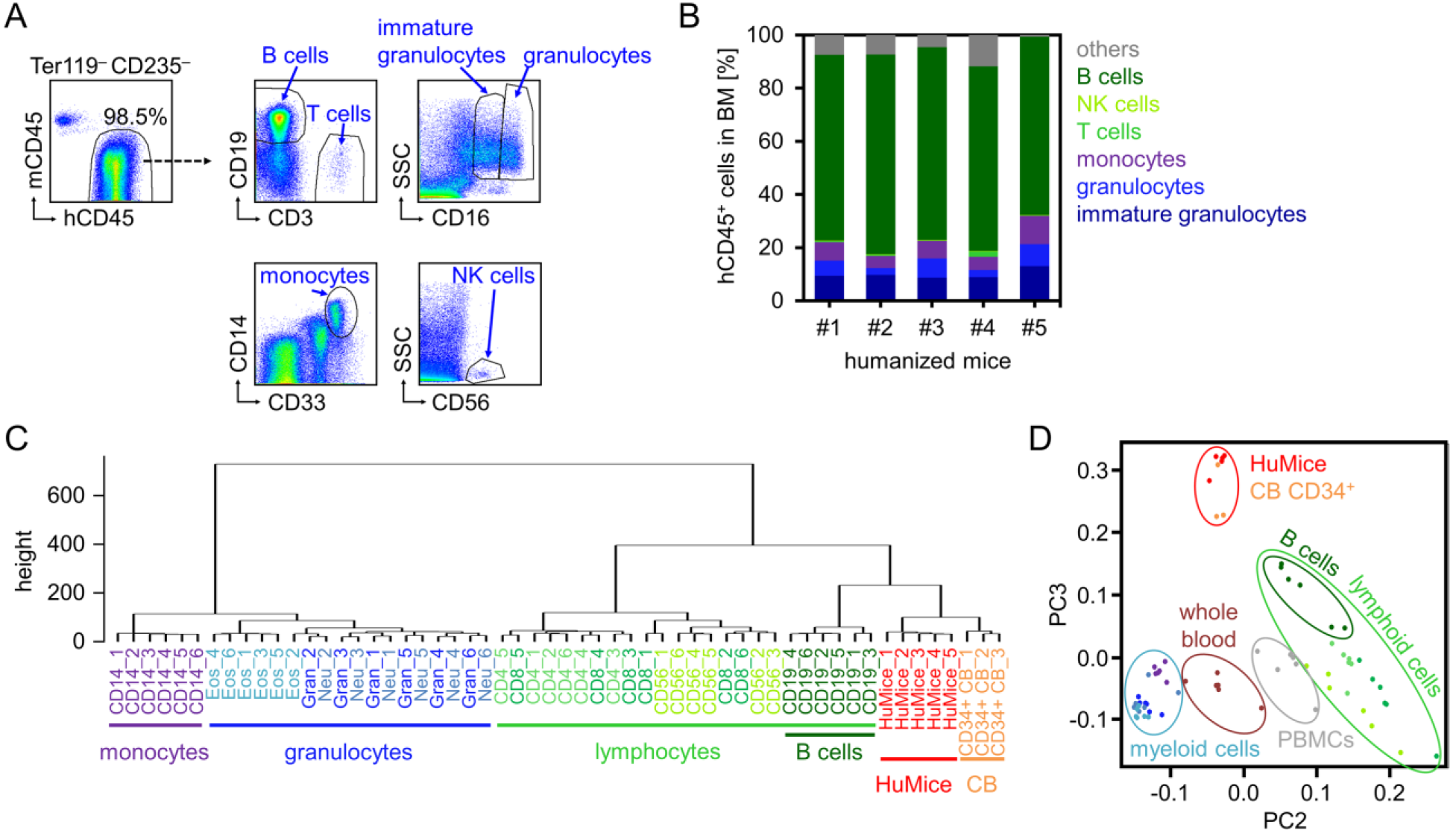
Phenotypic and epigenetic characterization of engrafted human hematopoietic cells. **(A)** Flow cytometric analysis of bone marrow (BM) 19 weeks after transplantation of human CD34^+^ cells into NSGW41 mice. Erythroid cells (Ter119^+^ or CD235^+^) were excluded and human CD45^+^ (hCD45^+^) cells were analyzed for the expression of cell type-specific surface markers of B cells (CD19), T cells (CD3), monocytes (CD14), NK cells (CD56), and granulocytes (CD16). **(B)** Cellular composition of hCD45^+^ cells in BM of five humanized mice. Cells described as “others” include stem and progenitor cells, myeloid progenitors, and dendritic cells. **(C)** Unsupervised hierarchical clustering of global DNA methylation (DNAm) profiles of various hematopoietic cell types purified from peripheral blood (monocytes, granulocytes, and lymphocytes; GSE35069) or umbilical cord blood (CB; GSE40799) compared to those of hCD45 sorter purified cells from BM of humanized mice (HuMice). **(D)** Principle component analysis (PCA) of the same hematopoietic subsets described in Fig. 1C. PBMCs = peripheral blood mononuclear cells.

To gain further insights into epigenetic changes of stably engrafted hematopoietic cells, we filtered for CpG dinucleotides with significant DNAm changes in HuMice *versus* CD34^+^ CB samples (adjusted *P* value < 0.05): 9 867 and 804 CpGs were hypo- and hypermethylated, respectively (Fig. 2A). For functional classification, we focused particularly on genes with significantly differentially methylated CpGs in promoter regions: Gene ontology (GO) analysis revealed highly significant enrichment of DNAm changes in hematopoietic categories (Fig. 2B), indicating that DNAm changes upon engraftment in HuMice are particularly associated with hematopoiesis and immune response.

**Fig. 2.**
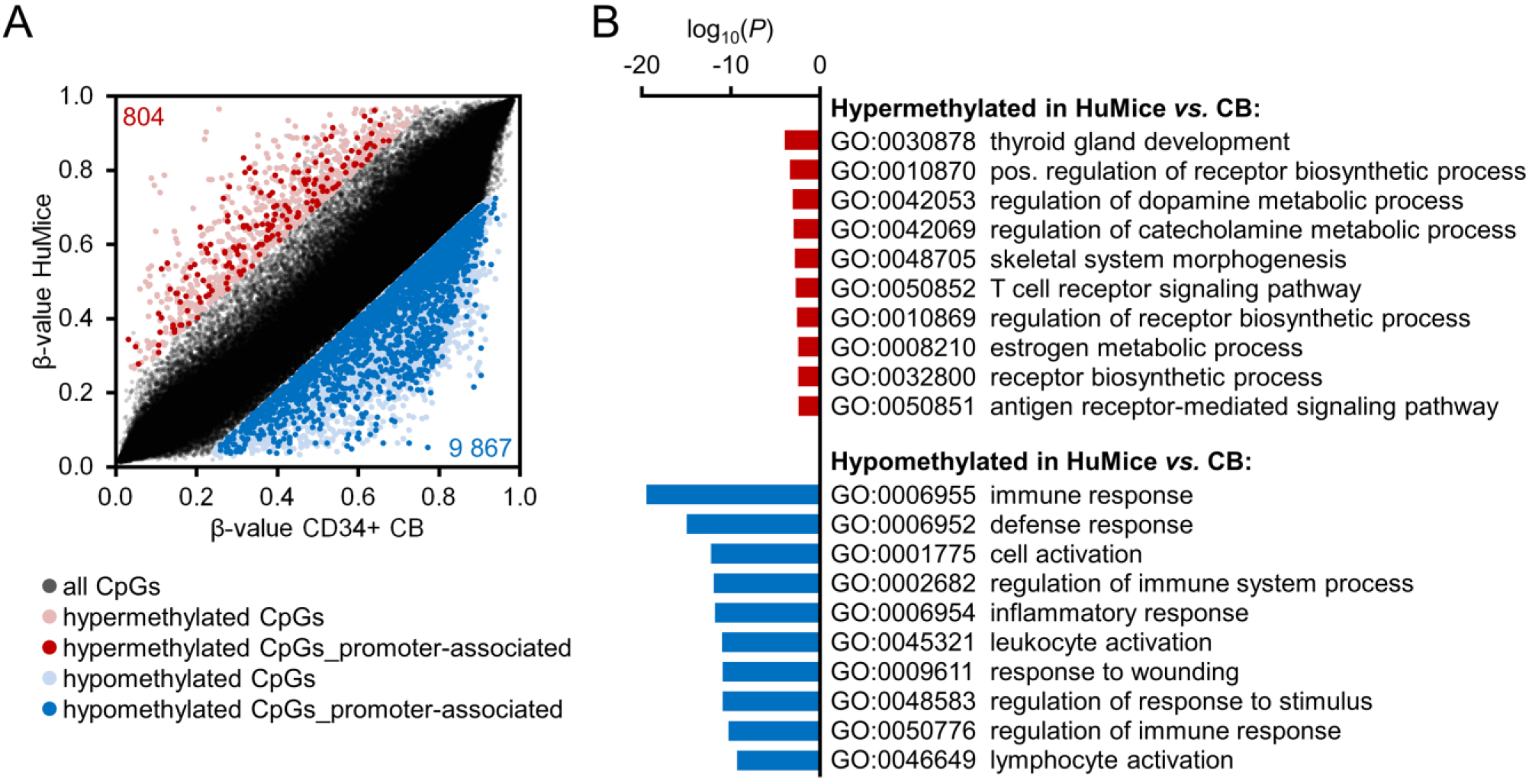
DNA methylation changes in human hematopoietic cells upon stable engraftment into mice. **(A)** Scatterplot of DNAm levels in humanized mice (HuMice) *versus* CD34^+^ cord blood (CB) samples. Significant hyper- and hypomethylation is highlighted in red and blue, respectively (delta of mean β-values > 0.2 or < −0.2; adjusted limma *P* value < 0.05). CpG sites that are associated with promoter regions (located in the 5’ untranslated region (5’UTR), 200 bp (TSS200), and 1 500 bp (TSS1500) upstream of transcription start site) (Sandoval et al., 2011) are more likely to be reflected in differential gene expression and were therefore highlighted in bold (2 425 CpGs and 169 CpGs, respectively). **(B)** Gene Ontology (GO) analysis of genes associated with differentially methylated CpG sites in promoter regions (one-sided Fisher’s exact *P* value). The most significant categories are exemplarily depicted (categories comprising more than 1 000 genes were not considered and similar categories are only listed once).

The cellular composition of hematopoietic subsets can be estimated based on DNAm patterns by deconvolution algorithms (Frobel et al., 2017; Houseman et al., 2012). When we applied the algorithm of Houseman *et al.* on DNAm profiles of HuMice the estimated relative cell counts were overall in line with immunophenotypic assessment (Fig. 3A). Furthermore, consistent with the high CD19^+^ content of engrafted cells, the promoter region of *CD19* was hypomethylated in engrafted cells with a very similar DNAm pattern as observed in sorted B cell populations from whole blood (Fig. 3B). In analogy, such characteristic patterns of B cells were also reflected in many other genes of B cell development. These results demonstrate that lineage-specific epigenetic profiles of normal human hematopoietic differentiation, particularly those of the B cell lineage, are recapitulated upon transplantation into mice.

**Fig. 3.**
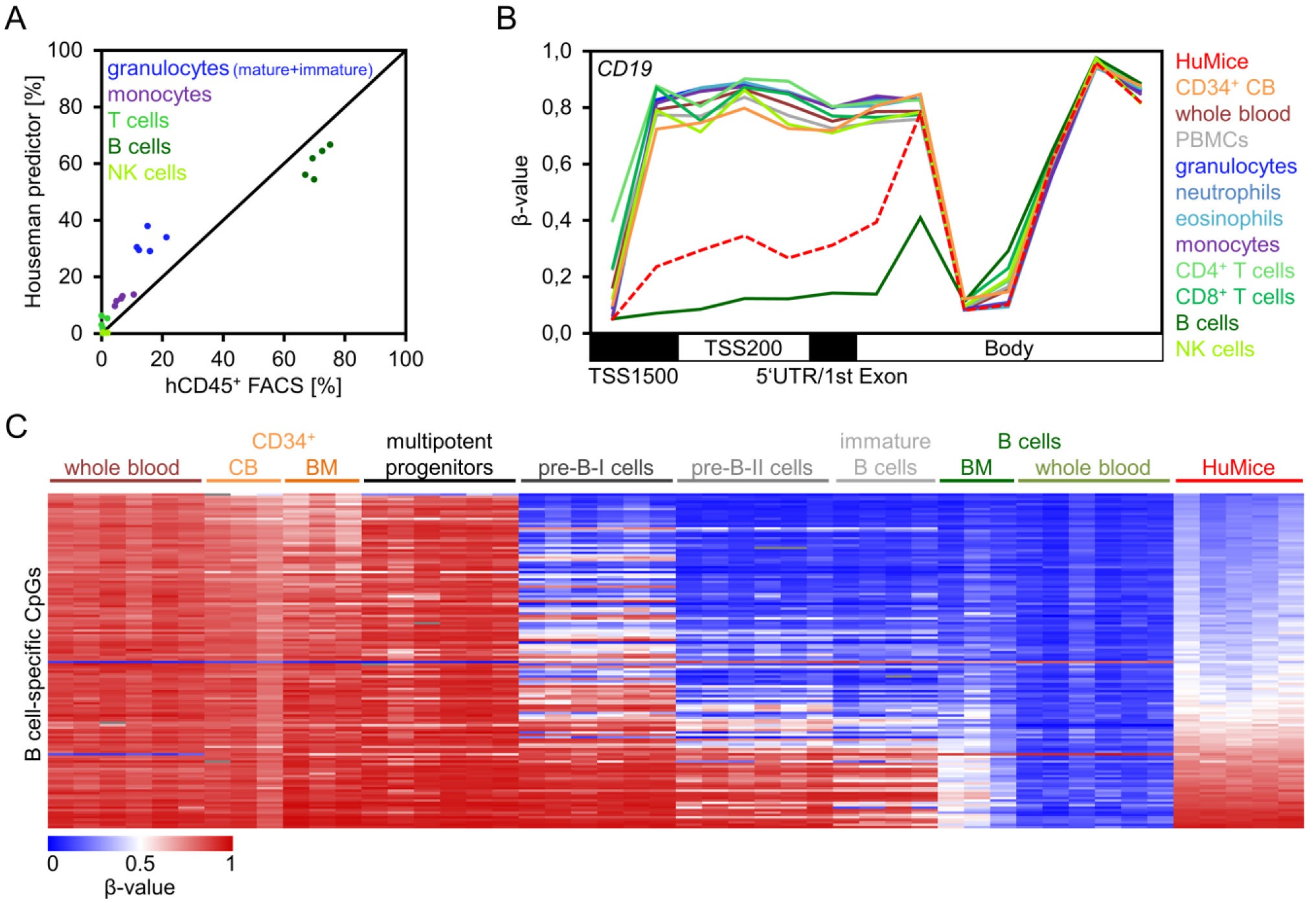
Lineage-specific DNA methylation changes. **(A)** Correlation of flow cytometric analysis of human CD45^+^ (hCD45^+^) bone marrow (BM) cells compared to results of a deconvolution algorithm to estimate the composition of cell types based on DNA methylation (DNAm) profiles (Houseman predictor) (Houseman et al., 2012). **(B)** DNAm levels (β-values) of CpGs within the gene *CD19* for various hematopoietic subsets from peripheral blood (GSE35069), umbilical cord blood (CB; GSE40799), and from BM of humanized mice (HuMice). **(C)** Heatmap of 330 CpGs with highest differential DNAm in B cells as compared to other mature cell types (GSE35069, delta of mean β-values > 0.7 or < −0.7; SD < 0.1). DNAm profiles of B cell precursors were subsequently included for comparison (GSE45459; multipotent progenitors = CD34^++^CD19^−^; Pre-B-I cells = CD34^+^CD19^+^; Pre-B-II cells = CD34^−^CD19^+^sIgM^−^; immature B cells = CD34^−^CD19^+^sIgM^+^; sIgM = surface IgM). The heatmap is sorted by mean DNAm levels in HuMice.

To better understand if DNAm patterns of normal human B cells are generally acquired in HuMice, we filtered for B cell-specific CpG sites (Frobel et al., 2017). The vast majority of these CpGs were hypomethylated in B cells and most of these were also hypomethylated in HuMice. In fact, the DNAm pattern of HuMice in these B cell-specific CpGs was perfectly in line with those of immature B cells in normal human development (GSE45459; isolated from fetal BM; Fig. 3C) (Lee et al., 2012). These findings further substantiate the notion, that the majority of engrafted cells resemble early B cell progenitor cells, which might contribute to their above mentioned close epigenetic relationship with CD34^+^ progenitor cells. On the other hand, the complex epigenetic modifications associated with lineage-specific differentiation are also initiated in the xenogeneic transplantation model.

It is generally anticipated that the microenvironment, the so called ‘stem cell niche’, has major impact on declining stem cell function in the elderly (Geiger et al., 2013). In fact, age-associated DNAm changes are acquired faster in short-lived mice than men (Stubbs et al., 2017; Wagner, 2017). With an average murine life-expectancy of about two years, the 19 weeks after transplantation might correspond to 15 years of human aging. To address the question if epigenetic aging is accelerated in the xenogeneic transplantation setting we initially focused on 99 age-associated CpGs that revealed high correlation with chronological age in blood (Weidner et al., 2014). In HuMice, most of these age-associated CpGs maintained the DNAm patterns of CB, even though some of them revealed moderate changes as observed upon aging (Fig. 4A). The corresponding age-predictor was not trained for CB samples and age-predictions were therefore underestimated for CB samples and HuMice (Fig. 4B). Alternatively, we used the age-predictors of Hannum *et al.* (Hannum et al., 2013) and Horvath (Horvath, 2013), and both models consistently indicated that 19 weeks after transplantation epigenetic aging is only moderately increased in mice (mean epigenetic-age increase of 6 and 0.7 years, respectively; Fig. 4C).

**Fig. 4.**
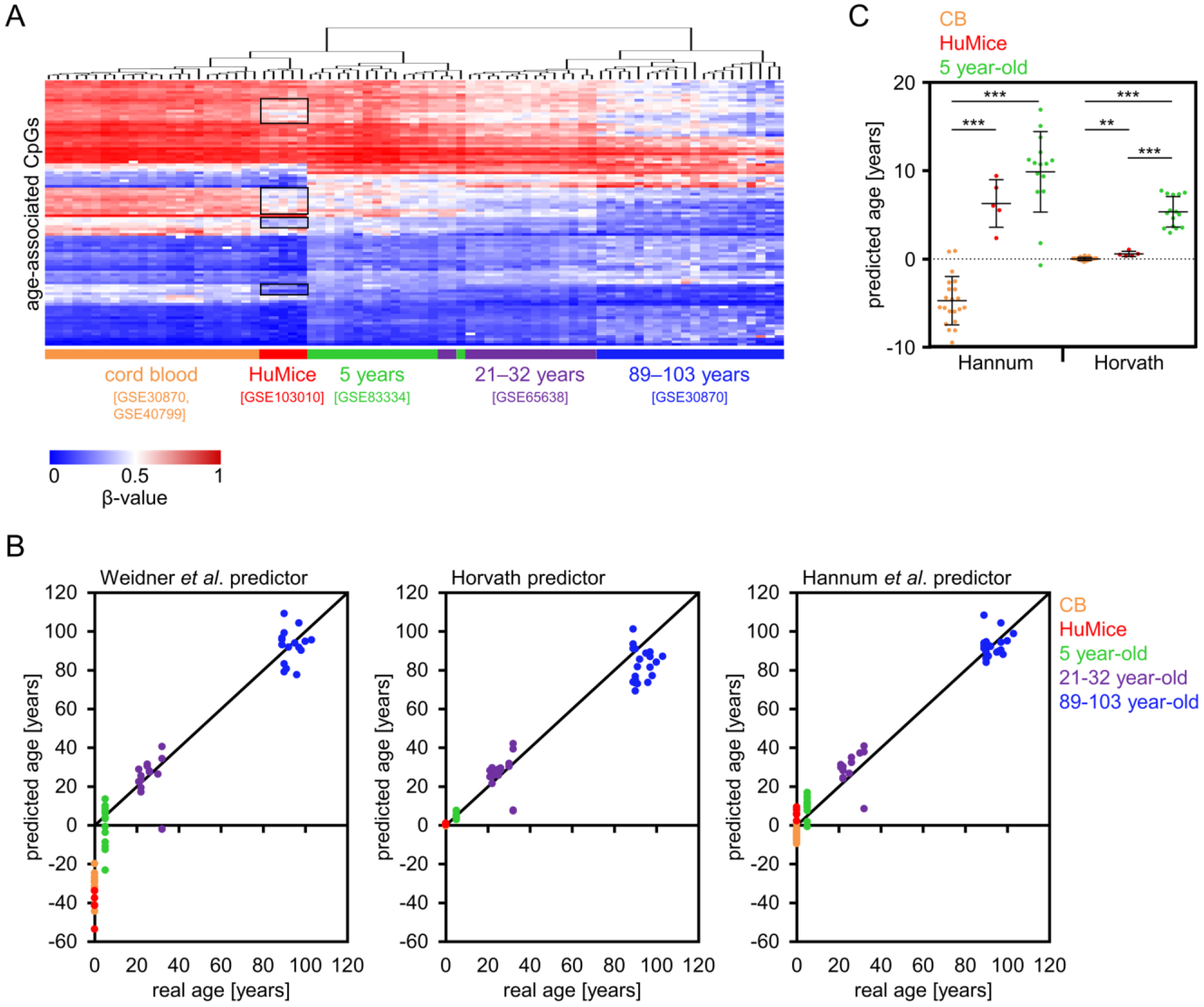
Epigenetic age predictions based on three different aging signatures. **(A)** The clustered heatmap represents DNAm of 99 age-associated CpG sites of the epigenetic age predictor of Weidner *et al.* (Weidner et al., 2014). Black rectangles highlight DNAm patterns of CpGs that may reveal moderate epigenetic age-acceleration in HuMice. **(B)** Three epigenetic aging predictors from Weidner *et al.* (Weidner et al., 2014), Horvath (Horvath, 2013), and Hannum *et al.* (Hannum et al., 2013) were used to predict donor age in 23 cord blood samples (CB; GSE40799, GSE30870), 15 whole blood samples of 5 year-old donors (GSE83334), 16 whole blood samples of 21-32 year-old donors (GSE65638), 20 peripheral blood mononuclear cell samples of 89-103 year-old donors (GSE30870), and the five samples from humanized mice (HuMice). The age predictor of Weidner *et al.* was not trained for CB and underestimates CB and HuMice samples to negative ages. **(C)** Epigenetic age predictors of Hannum *et al.* (Hannum et al., 2013) and Horvath (Horvath, 2013) were applied on DNAm profiles of CB cells, BM of HuMice, and peripheral blood of 5 year-old individuals. ** *P* < 0.01; *** *P* < 0.001 (two-tailed Student’s t-test).

Hematopoietic differentiation is governed by complex epigenetic mechanisms, which have to be triggered by the microenvironment. In this study, we demonstrate that the xenogeneic milieu of a murine transplantation model evokes very similar DNAm changes as observed in human hematopoiesis. On the other hand, the epigenetic makeup seems to be stalled on progenitor level. We have previously demonstrated that epigenetic age predictions in patients upon allogeneic hematopoietic stem cell transplantation correlate with donor age, while the microenvironment of elderly patients did not impose significant effects on age predictions (Weidner et al., 2015). This is in line with findings of this study, indicating that the epigenetic clock is hardly affected by a faster aging xenogeneic environment. Our study opens the perspective to investigate how the cellular microenvironment can be modified to facilitate better lineage-specific epigenetic maturation - to ultimately understand how hematopoietic cell-fate decisions are regulated.

## Materials and methods

### Humanized mice

Human hematopoietic stem and progenitor cells were isolated from human CB provided by the DKMS Cord Blood Bank Dresden. Three individual cord blood samples were pooled before Ficoll-Hypaque density centrifugation and magnetic enrichment for CD34^+^ cells according to the manufacturer’s instructions (Miltenyi Biotech) (Cosgun et al., 2014). 50 000 CD34^+^ cells were injected intravenously in 150 μl PBS / 5% FCS into five 7 to 9 week old unconditioned KIT-deficient NSGW41 mice. After transplantation mice were given neomycin-containing drinking water for 3 weeks.

### Flow cytometry and cell sorting

Nineteen weeks after transplantation, BM was collected from HuMice and prepared as described before (Cosgun et al., 2014). Leukocyte counts of murine (mCD45^+^, clone 30F11; eBioscience) or human (hCD45^+^, clone HI30; BioLegend) origin were determined. Furthermore, cells were stained using anti-human antibodies for CD3 (clone OKT3), CD19 (clone HIB19), CD33 (clone WM-53), CD235 (clone HIR2; all eBioscience), CD16 (clone 3G8; BioLegend), and CD14 (clone M5E2; BD Biosciences) and the anti-mouse antibody Ter119 (clone TER-119; eBioscience). Blocking reagent used was ChromPure mouse IgG (Jackson ImmunoResearch). Flow cytometric measurements were performed on a LSRII cytometer (BD Biosciences) and analyzed using FlowJo software (TreeStar). Sorting of human CD45^+^ cells was performed on a FACSAriaTM II (BD Biosciences).

### Analysis of DNA methylation profiles

Genomic DNA was isolated from 10^6^ sorted human CD45^+^ cells using the QIAmp DNA Blood Mini Kit (Qiagen) according to the manufacturer’s instructions including RNA digest. DNA isolated from BM of humanized mice was bisulfite converted with the EZ DNA Methylation Kit (Zymo Research) according to the manufacturer’s instructions. DNAm profiles were subsequently analyzed with the Infinium HumanMethylation450 BeadChip (Illumina). This platform features more than 450 000 cytosine guanine dinucleotides (CpG sites). DNAm levels at individual CpG sites are provided as β-values ranging from 0 (no methylation) to 1 (100% methylation).

### Bioinformatics

Our DNAm profiles of HuMice were compared with our previous data on human CD34^+^ CB cells (n = 3; GSE40799) (Weidner et al., 2013), B cells (n = 6; GSE35069) (Reinius et al., 2012), and various human hematopoietic subsets that were isolated from peripheral blood (granulocytes, neutrophils, eosinophils, CD4^+^ T cells, CD8^+^ T cells, NK cells, and monocytes; n = 6 per cell type; GSE35069) (Reinius et al., 2012). For comparative analysis of epigenetic age-predictions we used DNAm profiles of human CB cells (n = 20; GSE30870, and n = 3; GSE40799) (Simo-Riudalbas et al., 2015; Weidner et al., 2013), peripheral blood of 5 year-old individuals (n = 15; GSE83334) (Urdinguio et al., 2016), 21-32 year-old individuals (n = 16; GSE65638) (Xu et al., 2015), and 89-103 year-old individuals (n = 20; GSE30870) (Simo-Riudalbas et al., 2015). All of these DNAm profiles were generated on the same Infinium HumanMethylation450 BeadChip platform.

For further analysis, CpG sites located on the X and Y chromosomes were excluded, missing values were estimated by k-nearest-neighbor (kNN) imputation, and data was quantile normalized. Unsupervised hierarchical clustering according to Euclidean Distance and principle component analysis (PCA) were calculated in R. To estimate significant differences in DNAm we applied limma paired t-test in R (adjusted for multiple testing). *P* < 0.05 was considered as statistically significant. In addition, we selected for CpGs with a difference in mean β-values > 0.2 or < −0.2 in HuMice *versus* CD34^+^ CB samples to focus on CpGs with the highest difference. Functional classification of corresponding genes was performed with the GoMiner tool (Zeeberg et al., 2003). Enrichment of specific categories was calculated by the one-sided Fisher’s exact *P* value using all genes represented on the array as a reference. Epigenetic blood cell counts were calculated with the *Minfi* package in R using the *estimateCellCounts* function (Aryee et al., 2014; Houseman et al., 2012). For selection of B cell-specific CpGs we filtered for CpGs with a difference in mean β-value of > 0.7 or < −0.7 in B cells (GSE35069) compared to other mature hematopoietic cell types isolated from peripheral blood (granulocytes, neutrophils, eosinophils, CD4^+^ T cells, CD8^+^ T cells, NK cells, and monocytes; GSE35069). Furthermore, only CpG sites with a standard deviation (SD) < 0.1 within the respective cell type(s) were considered as described before (Frobel et al., 2017). Using this selection strategy, 330 B cell-specific CpG sites were identified. To estimate donor age based on DNAm, three different epigenetic age predictors were applied as described by Weidner *et al.* (Weidner et al., 2014), Hannum *et al.* (Hannum et al., 2013), and Horvath (Horvath, 2013). Statistical significance of deviations of predicted and chronological age was estimated with the two-tailed, unpaired Student’s t-test.

## Footnotes

### Author contributions

WW and JoF designed the study and JoF analyzed and formatted the data. SR and CW performed HuMice experiments, provided DNA samples, and analyzed FACS data. JuF supported bioinformatics. JoF and WW wrote the first draft of the manuscript and all authors read, edited, and approved the final manuscript.

### Ethics

All human samples were used in accordance with the guidelines approved by the Ethics Committee of the Dresden University of Technology. Animal experiments were performed in accordance with German animal welfare legislation and approved by the relevant authorities (Landesdirektion Dresden, Referat 24).

